# Proteomic changes induced by harmine in human brain organoids reveal signaling pathways related to neuroprotection

**DOI:** 10.1101/2021.06.16.448740

**Authors:** Karina Karmirian, Lívia Goto-Silva, Juliana Minardi Nascimento, Marcelo N. Costa, José Alexandre Salerno, Isis Moraes Ornelas, Bart Vanderborght, Daniel Martins-de-Souza, Stevens Rehen

## Abstract

Harmine is a β-carboline found in *Banisteriopsis caapi*, a constituent of ayahuasca brew. Ayahuasca is consumed as a beverage in native Americans’ sacred rituals and in religious ceremonies in Brazil. Throughout the years, the beneficial effects of ayahuasca to improve mental health and life quality have been reported, which propelled the investigation of its therapeutic potential to target neurological disorders such as depression and anxiety. Indeed, antidepressant effects of ayahuasca have been described, raising the question of which cellular mechanisms might underlie those effects. Previous animal studies describe potential neuroprotective mechanisms of harmine, including anti-inflammatory and antioxidant activities, and neurotrophin signaling activation. However, the cellular and molecular mechanisms modulated by harmine in human models remain less investigated. Here we analyzed the short-term changes in the proteome of human brain organoids treated with harmine using shotgun mass spectrometry. Harmine upregulates proteins related to synaptic vesicle cycle, cytoskeleton-dependent intracellular transport, cell cycle, glucose transporter-4 translocation, and neurotrophin signaling pathway. In addition, protein expression levels of Akt and phosphorylated CREB were increased after 24 hour-treatment. Our results shed light on the potential mechanisms that may underlie harmine-induced neuroprotective effects.

## Introduction

Harmine is a β-carboline, first isolated in the 1840s from *Peganum harmala* (Syrian rue) seeds, which are used as medicinal plants in Eastern countries^1,2^. In America, harmine is found in *Banisteriopsis caapi* (cipó-mariri) and is consumed in ayahuasca, a traditional psychotropic brew originated in the Amazon Forest by indigenous people. Since 1930, it has been used in religious ceremonies of ‘Santo Daime’ and ‘União do Vegetal’ churches, and later on, it became popular among people searching for its anecdotal beneficial effects in mental health^3,4^. Indeed, antidepressant effects induced by ayahuasca have been described in a randomized placebo-controlled trial targeting treatment-resistant depression^5^.

Ayahuasca is prepared by the decoction of *Banisteriopsis caapi* vines and *Psychotria viridis* leaves^6^, resulting in a beverage containing β-carbolines (harmine, harmaline, and tetrahydroharmine) and the psychedelic compound N,N-dimethyltryptamine (N,N-DMT) derived from *P. viridis*. The concentration of these components varies widely within preparations^7^. Therefore, it is of utmost relevance to investigate the mechanisms triggered by each compound, separately, in order to understand the mechanisms promoting beneficial neurological effects.

Harmine crosses the blood-brain barrier and binds to serotonin receptors, mainly to 5HT_2A_, followed by 5HT_2C_^8,9^ In addition, harmine inhibits monoamine oxidase A/B (MAO-A/B) and dual-specificity tyrosine-phosphorylation-regulated kinase 1A (DYRK1A)^10,11^. Since MAO-A/B catalyzes monoamine degradation, MAO inhibition elevates monoamine concentration in the synaptic cleft, which is one of the main mechanisms of action of antidepressant drugs^12^. DYRK1A is a major kinase associated with several functions during neurodevelopment, adult brain physiology, and neurodegenerative diseases^13-15^. Interestingly, DYRK1A inhibition has been described as a potential target for tauopathies and associated neurodegenerative diseases^16^.

Considering its molecular targets, harmine became a potential drug candidate for targeting neurological disorders or displaying neuroprotective effects. Indeed, several behavioral studies describe improvements in cognitive function mediated by acute or chronic treatment with harmine in mice models^17-21^. On a cellular level, harmine was found to stimulate the proliferation of neural progenitor cells, suggesting an effect in neurogenesis^22^. Further neuroprotective effects were also described in traumatic brain-injured mice, in which harmine treatment reduced the brain edema along with decreased levels of cell death and inflammatory markers in the hippocampus, leading to increased neuronal survival ^23^. Moreover, harmine displayed antioxidant effects, reducing lipid and protein oxidation along with increasing catalase and superoxide dismutase (SOD) activities in the prefrontal cortex and hippocampus of adult mice^24^. Besides the anti-inflammatory and antioxidant effects, harmine also modulates the neurotrophin signaling pathway by increasing brain-derived neurotrophic factor (BDNF) and phosphorylated TrkB protein levels in mice models^20,21,25^. Nonetheless, the effects of harmine in human neural cells remain less explored and investigating harmine effects in human models can bring new insights into harmine applications to treat neurological disorders.

Human brain organoids derived from induced pluripotent stem cells (iPSCs) exhibit structural and gene expression similarities with the human brain, recapitulating features of its cell diversity and cytoarchitecture^26-28^. Brain organoids have been used on neurological disease modeling, neurodevelopmental research, and drug screening^29-32^. Considering the wide range of applications, this model offers a great potential for translational research^33^. Here we analyzed proteomic alterations induced by short-term treatment with harmine in human brain organoids. We show that harmine alters proteins associated with heterocyclic compound binding, DNA replication, and antioxidant properties, which was expected considering the chemical structure of harmine and previous evidence supporting cell cycle regulation and antioxidant activity. Furthermore, we found that harmine modulates a wide range of processes including the synaptic vesicle cycle, protein folding, cytoskeleton-dependent transport, and glucose transporter-4 (GLUT4) translocation pathway. Harmine also upregulates AKT and increases p-CREB levels, indicating a positive regulation of the neurotrophin signaling pathway. Therefore, our findings help to confirm previously proposed mechanisms and describe novel pathways that may explain other potential effects triggered by harmine.

## Results

### Human brain organoids express harmine targets

First, we analyzed the basal expression of known harmine targets in human brain organoids at 45 days of formation (day-45). DYRK1A and 5HT_2A_ mRNA expression was confirmed with qualitative PCR for the presence of specific bands at 80 and 359 bp, respectively **(Fig. 1A)**. Then, DYRK1A and 5HT_2A_ protein expression were evaluated by immunocytochemistry (ICC) **(Fig. 1B, C)**. The pattern of DYRK1A staining was spread throughout the nuclei of brain organoid cells with notable labeling in proliferative zones, which was expected considering DYRK1A role in brain development **(Fig. 1B)**. The proliferative zones are rosette-like structures enriched with neural progenitor cells distributed radially and positive for nestin **(Fig. 2A)**. We also found 5HT_2A_ positive cells spread over the brain organoid. Some microtubule-associated protein 2 (MAP2) positive cells exhibit 5HT_2A_ staining, although 5HT_2A_ was also expressed in MAP2 negative cells, indicating that its expression is not restricted to neurons **(Fig. 1C)**.

**Figure 1:**
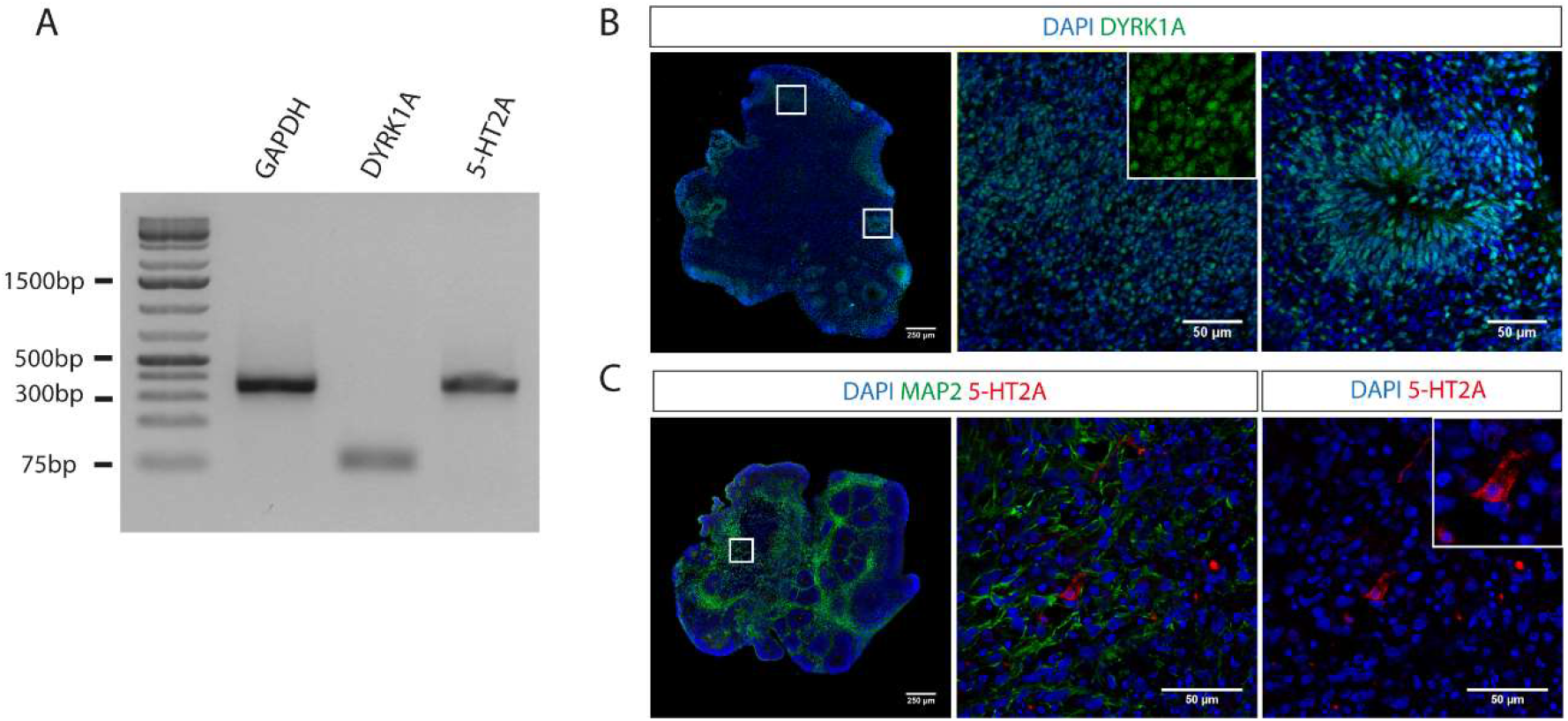
Expression of harmine targets in human brain organoids. **(A)** mRNA expression of DYRK1A and 5HT_2A_ evaluated by qualitative PCR in brain organoid at day-45. **(B)** Immunofluorescence image of brain organoid section at day-45 stained for DYRK1A (green). Note the proliferative zone in detail. **(C)** Immunofluorescence image of brain organoid section at day-45 stained for MAP2 (green) and 5HT_2A_ (red). Nuclei stained in blue (DAPI). Low magnification images: Scale bar = 250 μm; High magnification images: Scale bar = 50 μm.

**Figure 2:**
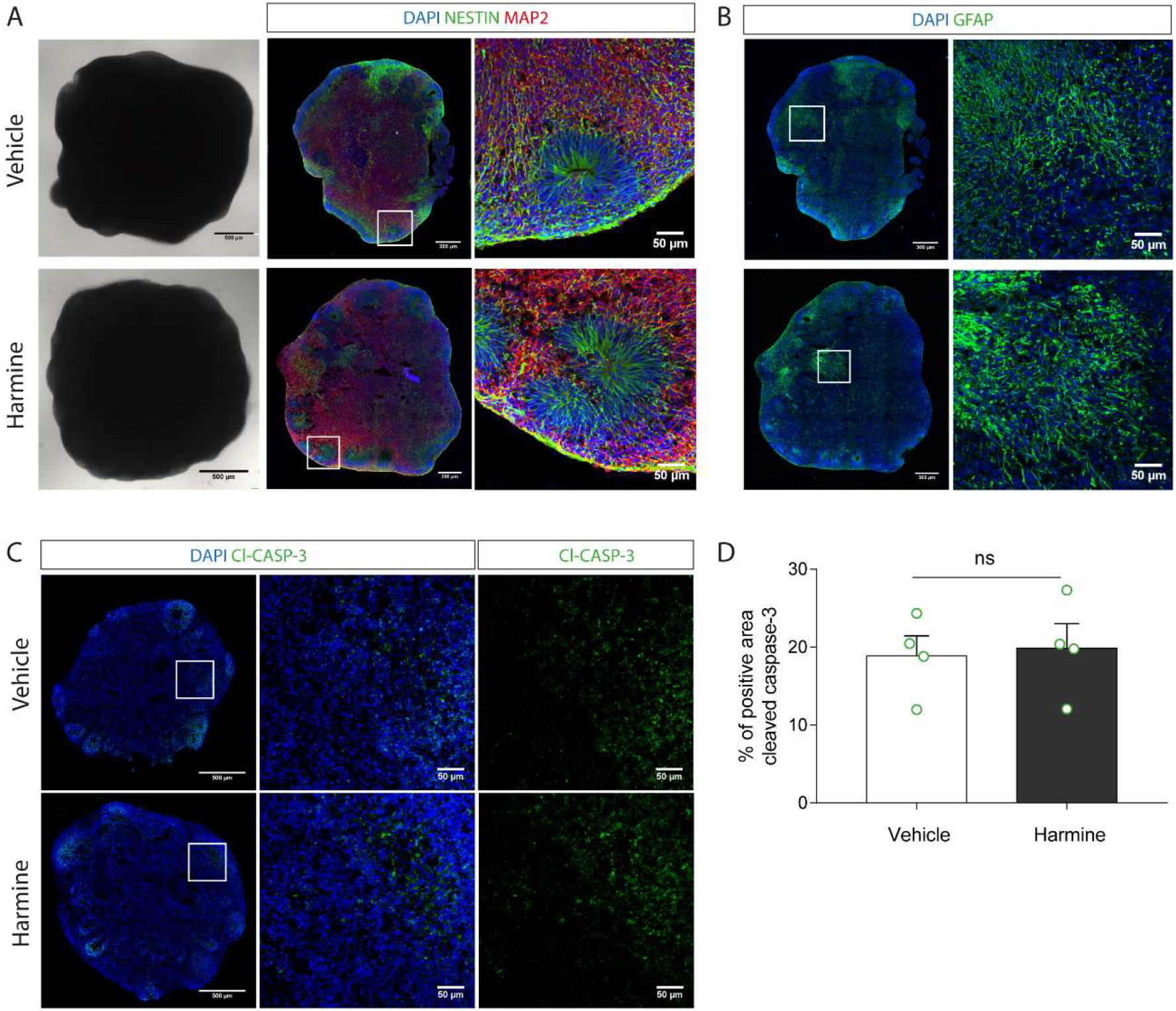
Morphological characterization of human brain organoids treated with harmine for 24 hours. **(A)** Brightfield image of brain organoids in culture obtained by digital inverted microscopy. Immunofluorescence images of brain organoid section (Day-45 - vehicle and harmine conditions) stained for Nestin (green) and MAP2 (red). Nuclei stained in blue (DAPI). **(B)** Immunofluorescence images of brain organoid section (Day-45 - vehicle and harmine conditions) stained for GFAP (green). Nuclei stained in blue (DAPI). **(C)** Immunofluorescence images of brain organoid section (Day-45 - vehicle and harmine conditions) stained for cleaved caspase-3 (green). Nuclei stained in blue (DAPI). **(D)** Quantification of the percentage of positive area stained for cleaved caspase 3. Each circle represents independent experiments. At least three sections of different brain organoids were quantified per batch. Low magnification images: Scale bar = 500 μm. High magnification images: Scale bar = 50 μm.

### Harmine does not affect brain organoid viability after 24 hours-treatment

Short-term effects induced by harmine have been demonstrated in mice models^21^. Therefore, we characterized brain organoids at day-45 treated with harmine for 24 hours. A previous study from our group described the increase of neural progenitor’s proliferation induced by harmine at 7.5 μM^22^. Most in vitro studies demonstrate harmine effects at a concentration range from 0.8 to 10 μM ^16,34^. Accordingly, we defined 7.5 μM harmine as the working concentration for our experiments. To evaluate whether harmine altered the cellular composition of brain organoids, we performed an ICC for neural markers including neural progenitors (nestin), mature neurons (MAP2), and glial cells (glial fibrillary acidic protein - GFAP) **(Fig. 2A, B)**. We observed that harmine did not induce significant change in the staining pattern of these markers, when compared to **controls (Fig. 2A, B)**. To ensure that 7.5 μM of harmine would not lead to any toxic effect, we performed an ICC for cleaved caspase-3, a well-known apoptosis marker **(Fig. 2C)**. After 24 hours, harmine did not induce any toxic effect, as quantified by the percentage of positive area for cleaved caspase-3 in **Fig. 2D**.

### Harmine induces proteome changes in heterocyclic compound binding, synaptic vesicle cycle, DNA replication, and microtubule-based processes

Changes in the proteome profile of brain organoids at day-45 treated with harmine were evaluated after a 24 hour-treatment. Proteins were identified using mass spectrometry based quantitative proteomics, as described in the experimental design workflow **(Fig. 3)**. After protein identification and data normalization, a total of 3695 proteins were identified at a false discovery rate of 1%. We found 155 proteins differentially expressed after harmine treatment (p-value cutoff = 0.05 among three independent batches), among which 93 proteins were up-regulated and 62 were down-regulated after short-term treatment with harmine.

**Figure 3:**
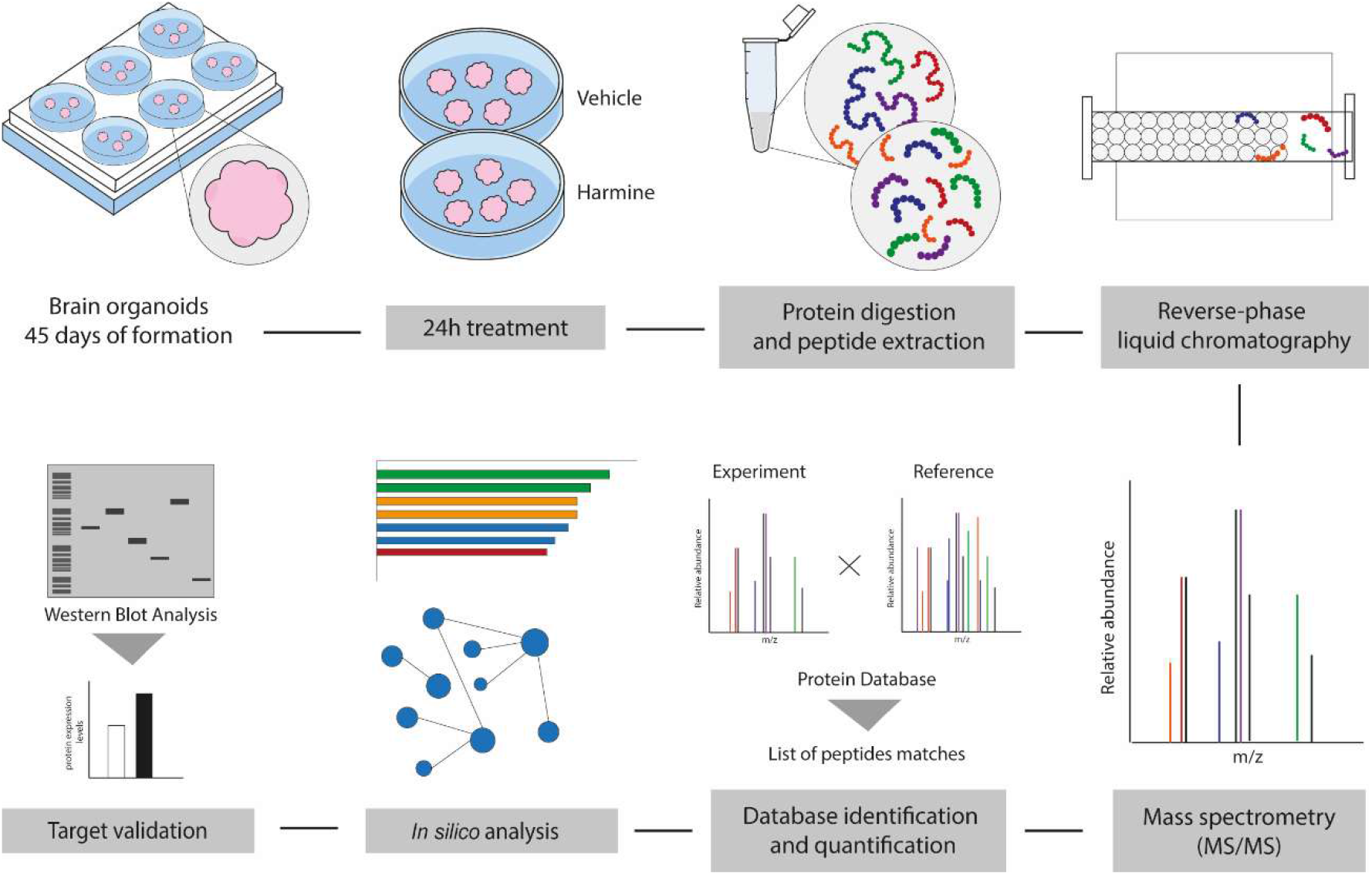
Experiment design workflow for proteomics. 45-day-old human brain organoids (five per batch) were treated with vehicle (DMSO) or harmine (7.5 μM) for 24 hours. Samples were analyzed using quantitative proteomics, based on twodimensional fractionation and high-resolution mass spectrometry. *In silico* analyses were performed to identify enriched biological processes and signaling pathways. Thus, some specific targets were validated by western blot analysis.

Functional enrichment analysis revealed several enriched processes using gene ontology (GO) databases for biological processes **(Fig. 4A)**, molecular function **(Fig. 4B)**, and cellular compartment annotations **(Fig. 4D)**. Among biological processes, cytoskeleton-dependent intracellular transport, vesicle trafficking, and microtubule-based processes were significantly enriched in harmine-treated organoids **(Fig. 4A)**. We performed an interactome analysis to better assess proteins related to these processes **(Fig. 4C)**. From that analysis, we found high and medium confidence interactions between kinesin light chain 1 (KLC1; fc = 5.42), kinesin-like protein KIF27 (fc = 1.79), trafficking kinesin-binding protein 2 (TRAK2; fc = 1.82), and kinesin-like protein KIF1B (fc = 0.73), all proteins related to axonal and microtubule transport. In addition, tubulin and microtubule-associated proteins were grouped into a second cluster related to microtubule-based processes. TUBB1 (fc = 1.68), TUBB2A (fc = 2.34), TUBA4A (fc = 3.79), MAP6 (fc = 1.66), tubulin-specific chaperone E (TBCE) (fc = 3.22) were up-regulated whereas tubulin-folding cofactor B (TBCB) were down-regulated (fc = 0.90).

**Figure 4:**
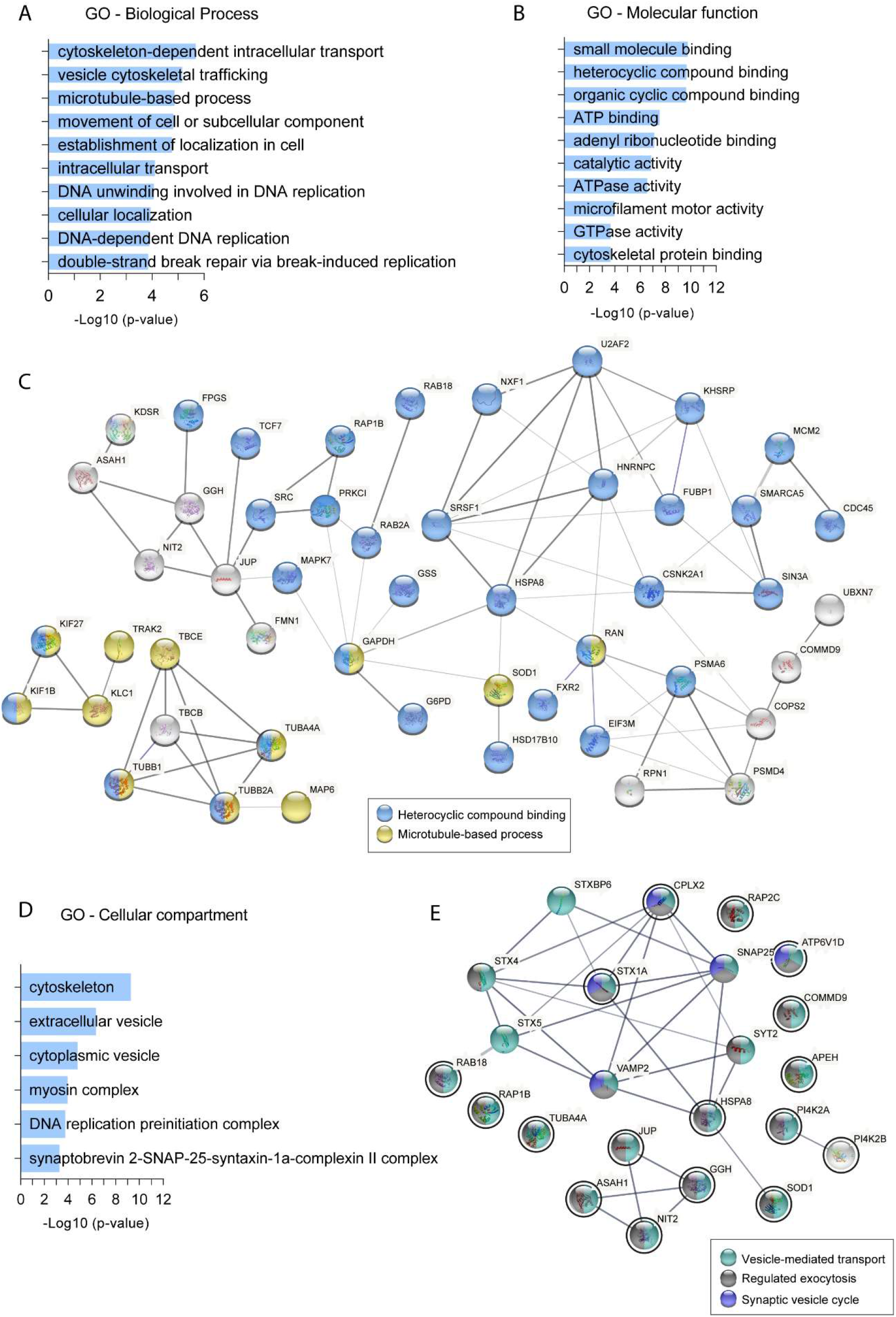
Functional enrichment analysis related to proteins differentially expressed after harmine treatment. **(A)** Biological process annotation, **(B)** Molecular function annotation, and **(C)**Interactome highlighting proteins associated with heterocyclic compound binding in light blue and microtubule-based process in yellow. Line thickness represents the interaction confidence, where high confidence interaction is represented with a thicker line. All proteins were identified in our dataset. Global protein-protein interaction (PPI) enrichment p-value = 0.0241. **(D)** Cellular compartment annotation and **(E)** Interactome highlighting proteins associated with vesicle-mediated transport in cyan, regulated exocytosis in gray, and synaptic vesicle cycle in dark blue. Proteins circled in black were identified in our dataset. Proteins in light gray were not annotated in those processes. Networks are based on the STRING database (https://string-db.org/).

Corroborating previous evidence supporting the role of harmine in the proliferation of human progenitor cells^22^, we also found DNA replication processes as enriched terms among biological process annotation **(Fig. 4A)**. Those processes comprise proteins such as DNA replication licensing factor MCM2 (fc = 1.12), ATP-dependent RNA helicase (RECQL, fc = 0.46), cell division control protein 45 (CDC45, fc = 0.50), DNA polymerase delta subunit 3 (POLD3, fc = 0.21), and DNA polymerase iota (POLI, fc = 2.51). MCM2 acts in MCM complex - together with other six MCM proteins - playing a role in DNA replication initiation and in the entry in S phase^35^. However, CDC45 and POLD3, both involved in DNA elongation and synthesis were down-regulated, suggesting that harmine displayed opposite effects in proteins related to cell cycle progression.

Interestingly, ATP binding and heterocyclic compound binding were highly enriched terms associated with molecular function annotation **(Fig. 4B)**. Noteworthy, harmine is an organic compound with heterocyclic structure that interacts with the ATP binding site of DYRK1A^10^. Therefore, it suggests that harmine activated pathways in human brain organoids follow well-described mechanisms. Furthermore, we analyzed the interaction of heterocyclic compound binding-related proteins, which may provide clues to other unknown mechanisms **(Fig. 4C)**. This subset of proteins formed a highly interactive cluster with different classes of proteins. We found three RAS-related proteins — RAP1B (fc = 6.28), RAB2A (fc = 0.74), RAB18 (fc = 2.30) — along with proto-oncogene tyrosine-protein kinase (SRC) (fc = 1.41) which participates in different intracellular signaling pathways, such as receptor tyrosine kinases, cytokine receptors, as well as G protein-coupled receptors. In fact, we found guanine nucleotide-binding protein G(q) subunit alpha (GNAQ) up-regulated in our dataset (fc = 3.59). Proteins related to mRNA processing (KHSRP, fc = 1.66; U2AF2, fc = 4.83; SRSF1, fc = 0.32), mRNA translation (EIF3M, fc = 5.01), cell cycle (MCM2, fc = 1.12; CDC45, fc = 0.50), and metabolism (G6PD, fc = 0.53) were also associated to heterocyclic compound binding. In addition, the antioxidant enzymes superoxide dismutase (SOD1, fc = 1.32; annotated in microtubule-based process) and glutathione synthetase (GSS, fc = 2.68; annotated in heterocyclic compound binding) were up-regulated, a possible correlation to harmine antioxidant effects as previously demonstrated^24^.

Considering cellular compartment annotation, extracellular vesicle and cytoskeleton represented the most enriched terms **(Fig. 4D)**. However, two proteins annotated in the SNARE complex (STX1A, fc = 1.18; and CLPX2, fc = 3.00) were upregulated, leading to a fold enrichment value of 88.01 and p-value of 5.53E-04 of that complex. We then analyzed the contribution of those two upregulated proteins (STX1 and CLPX2) and their connection to other proteins related to the synaptic vesicle cycle. The interactome included other proteins from significantly enriched processes, such as regulated exocytosis and vesicle-mediated transport. **(Fig. 4E)**. We found high confidence interactions (0.7) between STX1A and CLPX2 with other proteins associated with synaptic vesicle fusion, such as VAMP2 and synaptosomal associated protein 25 (SNAP25), which were not identified in our dataset. In addition, many proteins share the annotation of regulated exocytosis and vesicle-mediated transport such as RAS-related proteins (RAP1B, RAP2C, and RAB18) and phosphatidylinositol 4-kinase type 2-alpha (PI4K2A; fc = 3.30).

### Harmine regulates pathways related to cell cycle, protein folding, and vesicle-mediated transport

To analyze additional pathways that respond to harmine treatment, we performed an unbiased overrepresentation analysis on the Reactome database at p-value cut-off ≤ 0.05. Among the 69 enriched pathways, we found 4 higher hierarchy events corresponding to G2/M checkpoints (FDR = 7.23E-1; p-value = 2.33E-2), DNA replication (FDR = 5.12E-1; p-value = 1.35E-2), protein folding (FDR = 1.32E-1; p-value = 5.36E-4), and vesicle-mediated transport, including GAP junction trafficking and regulation (FDR = 4.96E-1; p-value = 9.72E-3) and translocation of glucose transporter type 4 (GLUT-4) to the plasma membrane (FDR = 7.5E-1; p-value = 3.31E-2) **(Fig. 5A)**.

**Figure 5:**
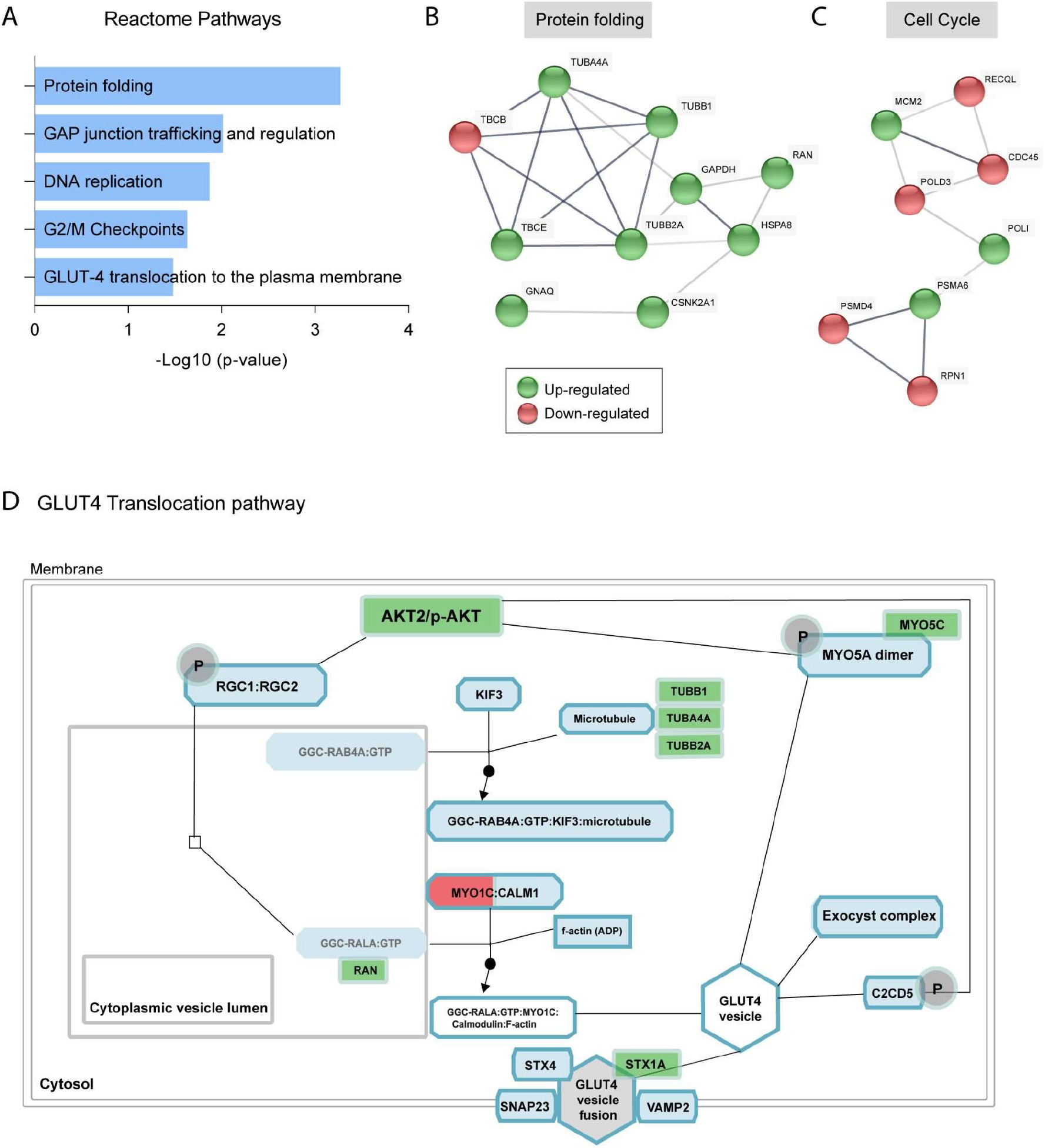
Overrepresentation analysis of enriched pathways induced by harmine treatment. **(A)** Enriched pathways among differentially regulated proteins. **(B)** Interactome of proteins annotated in protein folding pathway. **(C)** Interactome of proteins annotated in DNA replication and G2/M checkpoint grouped as cell cycle. Line thickness represents the interaction confidence, where high confidence interaction is represented with a thicker line. **(D)** Schematic representation of GLUT4 translocation pathway (FDR = 7.5E-1; p-value =3.31E-2). Components colored in green represent upregulated proteins and red downregulated proteins. Signaling components colored in blue were not found in our dataset. Overrepresentation analysis was performed with Reactome (https://reactome.org/).

The protein folding pathway comprises two enriched processes related to chaperonin-mediated protein folding (p-value = 4.73E-3) and post-chaperonin tubulin folding pathway (p-value = 7.38E-6). Among 10 differentially regulated proteins annotated in the protein folding pathway, we highlight the up-regulation of TBCE and down-regulation of TBCB, as represented in the interactome in **Fig. 5B**. These proteins actively interact during the regulation of tubulin heterodimer dissociation and further organization of the microtubule cytoskeleton^36^. While TBCE is required for proper maintenance of the neuronal network, evidence suggests that decreased TBCB levels induce axonal growth^37^. Therefore, the regulation of those proteins might be intimately related to the neuronal network organization in brain organoids.

Corroborating the GO annotations of biological process and cellular compartment **(Fig. 4A, D)**, the DNA replication pathway was significantly regulated, including specific pathways related to assembly (p-value = 3.39E-3) and activation (p-value = 3.62E-2) of the pre-replicative complex and synthesis of DNA (p-value = 6.28E-3). The synthesis of DNA pathway also comprises DNA strand elongation (p-value = 9.72E-3) and unwinding of DNA (p-value = 1.38E-2). Those pathways share the annotation of key proteins with the G2/M checkpoint process, which both indicate a regulation of cell cycle progression. However, as aforementioned, distinct stages of the cell cycle were differentially regulated considering up- and down-regulation of key proteins such as MCM2, POLD3, and CDC45 **(Fig. 5C)**. In addition, components of proteasome complexes, such as PSMA6 (fc = 1.58) and PSMD4 (fc = 0.79) were also annotated in DNA replication and G2/M checkpoint pathways. Interestingly, such proteins were also differentially regulated, which could possibly indicate distinct roles of protein degradation involved in cell cycle, apoptosis, and DNA repair.

Considering the vesicle-mediated transport pathways, we found two pathways that were not previously annotated in GO analysis. GAP junction trafficking pathway include tubulin proteins (TUBB1, TUBA4A, and TUBB2A), SRC, and clathrin light chain A (CLTA; fc = 0.67), the latter the only down-regulated protein, which is required for gap junction internalization^38^ Moreover, the translocation of GLUT-4 to the plasma membrane was also significantly enriched within vesicle-mediated transport pathways. Considering the growing evidence on harmine’s role as an anti-diabetic drug, the regulation of this pathway shed light to a possibly novel mechanism in which harmine may regulate brain insulin resistance and promote cognitive benefits^39^.

GLUT-4 translocation pathway includes 8 differentially regulated proteins that we highlighted in **Fig. 5D**. Following the schematic representation of the signaling pathway, RAB4A/GTP activates KIF3, inducing the formation of a complex with microtubules. Although KIF3 was not identified in proteomics, this signaling component was significantly enriched considering other kinesins found in the dataset. Microtubule components were upregulated, such as TUBB1, TUBB2A, and TUBA4A. Another positively regulated protein was AKT2 (fc = 2.46) that phosphorylates RGC1/RGC2, which participates in RAS-related protein Ral-A (RALA) hydrolysis of GTP. In this step, another upregulated protein found was GTP-binding nuclear protein Ran (RAN, fc = 1.68), a GTPase with positive interaction in this process. AKT2 can also phosphorylate MYO5A, which participates in GLUT4 vesicle formation. MYO5C upregulation (fc = 2.17) indicates a possible interactor in this step. The only downregulated protein associated with the GLUT4 translocation process was MYOC1 (fc = 0.073), which participates in a complex with calmodulin. After GLUT4 vesicle formation, fusion to the plasma membrane is mediated by VAMP2, SNAP23, and STX4^40^ **(Fig. 5)**. Those proteins interact with STX1A, which was upregulated (fc = 1.18) after harmine treatment.

### Harmine modulates proteins related to neurotrophin signaling pathway

Considering previous evidence showing increased BDNF levels after acute and chronic treatment with harmine in mice hippocampus^20,21^, along with p-Trk-B upregulation^25^, we decided to investigate proteins potentially related to neurotrophin signaling pathway in the proteomic data set. Although the neurotrophin signaling pathway was not significantly enriched, we found three relevant proteins upregulated as colored in green in **Fig. 6A**. SRC was 1.41-fold increase in harmine-treated organoids, along with RAP1B (fc = 6.28) and AKT2 (fc = 2.46). This could predict a possible positive regulation of targets downstream of the neurotrophin binding to the Trk receptors, which could be also activated by neurotrophin increase itself.

**Figure 6:**
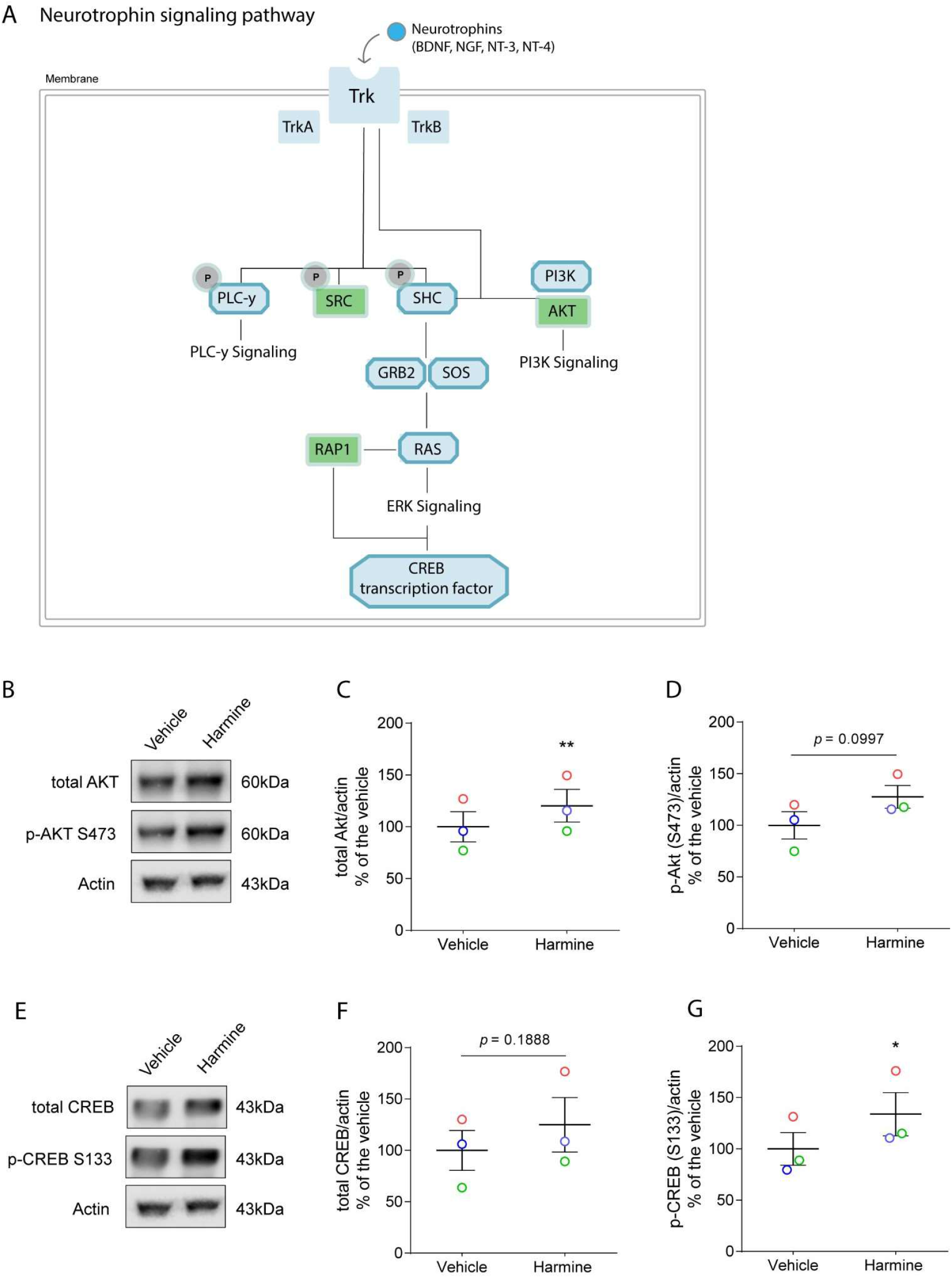
Effects of harmine on neurotrophin signaling pathway. **(A)** Schematic representation of main proteins involved in neurotrophin signaling pathway. Proteins colored in green were found upregulated in the proteomic dataset. **(B)** Western blot analysis for total AKT and p-AKT S473 in brain organoids treated with harmine for 24 hours. **(C)** Quantification of total AKT **(D)** and p-AKT protein expression by densitometry normalized by actin. **(E)** Western blot analysis for total CREB and p-CREB S133 in brain organoids treated with harmine for 24 hours. **(F)** Quantification of total CREB **(G)** and p-CREB protein expression by densitometry normalized by actin. Each circle with paired colors represents independent experiments. Protein extraction was performed with at least five organoids per batch. **p = 0.034. * p = 0.0257.

To test whether neurotrophins were also modulated by harmine treatment, we performed RT-qPCRs in brain organoids treated with harmine or vehicle for 24 hours **(Supp Fig 1)**. Among three batches analyzed, we found a trend of increase — not statistically significant - in BDNF, nerve growth factor (NGF), and neurotrophin 3 (NT-3) mRNA expression **(Supp Fig 1A-C)**, while not for neurotrophin 4 (NT-4) **(Supp Fig 1D)**.

Although we did not find a significant increase in neurotrophins mRNA levels, the proteins upregulated in proteomics - AKT, RAP1, and SRC - could predict a positive regulation of the neurotrophin signaling pathway. Indeed, we confirmed AKT upregulation in brain organoids treated for 24 hours by western blot analysis **(Fig. 6B, C)**. Total AKT was significantly increased (1.20-fold) in brain organoids from three independent batches **(Fig. 6C)**. The phosphorylated form of AKT exhibits an increasing trend although not statistically significant **(Fig. 6D)**. Furthermore, we then decided to evaluate the phosphorylation of downstream targets of those proteins such as cAMP-response element binding protein (CREB) **(Fig. 6E-G)**. CREB phosphorylated form (S133) had a 1.34-fold increase in brain organoids 24 hours-treated with harmine **(Fig. 6G)** whereas total CREB exhibits a trend not statistically significant **(Fig. 6F)**. Taken together, these data suggest that neurotrophin signaling pathway might be positively regulated by short-term harmine treatment.

## Discussion

Harmine is a β-carboline alkaloid with binding properties to serotonin receptors and a potent inhibitor of DYRK1A and MAO-A/B. Here we characterized the expression of harmine’ targets - 5HT_2A_ and DYRK1A - indicating the potential responsiveness of brain organoids to harmine. Next, we treated human brain organoids with 7.5 μM harmine for 24 hours and performed a proteomic analysis. Functional enrichment analysis revealed that harmine modulates proteins related to a wide range of processes, including synaptic vesicle cycle and neurotrophin signaling pathway, indicating a potential regulation of pathways associated with neuroprotection. Moreover, harmine regulates microtubule-based processes, including protein folding and vesicle-mediated transport.

Harmine induced changes in the level of proteins related to ATP binding and heterocyclic compound binding processes. This is in line with harmine being a heterocyclic compound. As such, harmine interacts with residues from the ATP binding site, including that of DYRK1A. This leads to ATP displacement and therefore inhibits phosphorylation of DYRK1A substrates^10^. DYRK1A interacts with several targets, including serine/arginine-rich splicing factor 1 (SRSF1) and RAS-related proteins that were both regulated by harmine and annotated in heterocyclic compound binding term^41,42^. Regarding other harmine potential targets, G(q) subunit alpha was increased by 3.59-fold, suggesting an interaction of harmine with G-protein coupled receptors, possibly including the 5HT_2A_ receptor. Although harmine binds to 5HT_2A_ receptor (Ki = 230-397 nM)^9^, it is unknown whether harmine acts as an agonist or antagonist. A single piece of evidence suggests an agonist mechanism due to increased dopamine efflux after an evoked potential in murine cerebral tissue treated with harmine, which was reversed by ketanserin (selective antagonist of 5HT_2A_)^43^.

Harmine also modulated proteins associated with DNA replication, although it displayed opposite effects on key proteins associated with cell cycle progression, such as the upregulation of MCM2 and downregulation of CDC45, both components of the Cdc45-MCM-GINS complex required for DNA replication in the S phase^44^. Although it is still unknown the direct impact of harmine-mediated DYRK1A inhibition in MCM2 and CDC45 expression, the role of DYRK1A on the regulation of cyclin-dependent kinase inhibitor 1B (Kip1) is well-understood. DYRK1A loss-of-function leads to Kip1 down-regulation, and therefore inducing cell proliferation. On the other hand, DYRK1A overexpression leads to cell cycle arrest due to Kip1 up-regulation^45^. Thus, DYRK1A expression is temporally regulated during neural progenitor proliferation and differentiation^14^, since *knockout* or overexpression models of DYRK1A share similar outcomes of brain development defects and dysregulation of neuronal differentiation^46^. This evidence emphasizes DYRK1A as a dosage-sensitive gene according to each step of neural development. Harmine induces the proliferation of neural progenitor cells *in vitro* after 4-days through DYRK1A inhibition^22^. Considering the transient expression of DYRK1A during neural proliferation and differentiation^14^, harmine may affect cells at distinct stages of development as well, as seems to be the case of those present in brain organoids.

Antioxidant properties of harmine have been described, including increased activity of catalase and SOD1 in the mouse prefrontal cortex and hippocampus induced by acute or chronic harmine treatment, along with a decrease in lipid and protein oxidation^20,24^. We observed that SOD1 and GSS were upregulated in harmine-treated organoids, supporting antioxidant activity in this model. SOD1 is the cytoplasmic isoform of superoxide dismutase and acts on redox cycle producing H_2_O_2_, which will be further reduced in H_2_O by catalase and glutathione peroxidases (GPx)^47^. On the other hand, GSS acts in glutathione biosynthesis, catalyzing the condensation of γ-glutamylcysteine and glycine to glutathione^48^. Therefore, both enzymes act in distinct stages contributing to the reduction of reactive oxygen species. Oxidative stress is a harmful state closely related to the pathophysiology of depression, as well as in neurodegenerative diseases, contributing to neuronal degeneration and synaptic dysfunction^49-51^. The antioxidant activity elicited by harmine is in accordance with its antidepressant effects demonstrated in mice models, which also correlates to antioxidant properties of classic antidepressant drugs such as fluoxetine and imipramine^20,24,52^. Furthermore, the scavenging effect of harmine was assessed using 2-deoxy-D-ribose degradation assay, in which harmine and other β-carbolines reduced 2-deoxy-D-ribose degradation induced by Fe+3, H202, EDTA, and ascorbate^53^. Additionally, β-carbolines exerts protective effects on dopamine- and 6-OHDA-induced alterations in synaptosomes and brain mitochondria, through the inhibition of thiol oxidation^53^.

Cytoskeleton-dependent intracellular transport was the most enriched among biological processes modulated by harmine. We were able to identify members of kinesin and dynein classes differentially expressed in the proteomic dataset, including KLC1, KIF27, KIF1B, TRAK2, dynein heavy chain 14, and dynein regulatory complex subunit 4. Axonal trafficking of neurotrophins contributes to neuronal survival and neuroprotection and its disruption implicates several neurological disorders^54^. Neurotrophin binding at axonal termini stimulates retrograde axonal trafficking^55^. We observed that proteins related to the neurotrophin pathway were up-regulated in our dataset, which could explain the regulation of axonal trafficking proteins by harmine treatment. Phospho-CREB was activated by harmine after 24-hours, and together with SRC, AKT and RAP1, these results suggest that harmine activates TrkB signaling, corroborating previous data on p-TrkB increased expression upon harmine injection in mice^25^. CREB is one of the final targets of the TrkB pathway and its activation results in transcription required for synaptic plasticity and memory consolidation^56^.

Synaptic vesicle cycle was enriched in proteomic dataset due to the upregulation of syntaxin 1A (STX1A) and complexin 2 (CPLX2). Indeed, STX1A and CPLX2 closely interact with VAMP2 and SNAP25, both components of SNARE complex, which mediates the fusion of synaptic vesicles^57^. Recently, Liu et al demonstrated the role of harmine in GABAergic transmission (but not glutamatergic) in basal amygdala projection neurons. Harmine enhanced the release of GABA from the presynaptic terminal^58^. Moreover, CPLX is involved in neurotransmitter release from both excitatory and inhibitory synapses^59^. These results suggest that harmine may regulate synaptic vesicle cycle by regulating the expression of STX1A and CPLX2.

Members of the SNARE complex can be related not only to synaptic vesicles but also to the fusion of vesicles associated with the translocation of protein to cell surface^60^. In fact, the signaling pathway linked to GLUT-4 translocation was significantly enriched in brain organoids after harmine treatment. GLUT-4 is an insulin-regulated glucose transporter that plays a crucial role in the homeostasis of glucose through its uptake from the circulation^61^. Although GLUT-1 and GLUT-3 are highly expressed in the brain, the translocation of GLUT-4 mediated by insulin was already demonstrated in the rat hippocampus, along with the activation of PI3K/AKT^62^. GLUT-4 translocates to the cell membrane after memory training tasks and GLUT-4 inhibition leads to impairment of long-term memory formation along with decrease in the levels of BDNF^63^. Therefore, harmine-mediated GLUT-4 translocation could possibly trigger a mechanism with potential benefits in cognitive function and brain insulin resistance^64^. Nevertheless, further studies are needed to confirm the role of harmine in the regulation of GLUT-4 translocation to the plasma membrane, as suggested by our data.

Diabetes has been strongly correlated to cognitive dysfunction, which led to the investigation of the protective effect of harmine in diabetic rats^25^. Harmine alleviated the impairment of learning and memory caused by diabetes. The authors also observed that harmine inhibited the formation of inflammasomes, along with a reduction in the levels of cleaved caspase 1, IL-1β, and IL-18. In addition, BDNF/TrkB signaling was significantly activated by harmine^25^. In fact, harmine stimulates BDNF production/signaling, although the underlying mechanisms are not well understood^20,21,25^. BDNF, NGF and NT-3 mRNA expression had a trend to increase in brain organoids at 24 hours of treatment. From our current data, it cannot be discarded that the effects of harmine may involve sustained signaling from neurotrophins with a decrease in signal degradation rather than an impact in transcript levels. Therefore, more experiments are needed to investigate the regulatory mechanisms of harmine in neurotrophin signaling.

In the present work, we demonstrate short-term effects of harmine in human brain organoids using proteomics. Among several pathways analyzed, harmine upregulated proteins related to synaptic vesicle cycle, antioxidant properties, axonal transport, and neurotrophin signaling pathway. Proteomics revealed an enriched pathway related to GLUT-4 translocation to the plasma membrane. It is noteworthy that deficits in synaptic function, insulin and neurotrophin pathways are commonly associated with neurological disorders, such as depression and neurodegenerative diseases, including Alzheimer’s disease^65–70^. Therefore, our results suggest novel cellular mechanisms on which harmine may exert metabolic and cognitive effects in the central nervous system that might underlie therapeutic properties still unexplored.

## Materials and Methods

### Brain organoid formation

Control iPSC line (GM23279A) from the NIGMS Human Genetic Cell Repository was obtained commercially from Coriell Institute. iPSCs were maintained with mTeSR1 medium and upon confluence passaged manually or with EDTA. High quality colonies with no differentiated cells were used for brain organoid formation following a previously described protocol^71^.

Briefly, human iPSC cells were detached using Accutase for 5min at 37 °C and followed dissociated into single cells with phosphate-buffered saline (PBS; LGC Biotechnology, USA) containing 10μM ROCK inhibitor (ROCKi, Y27632; Merck Millipore, USA). Cells were centrifuged at 300 rpm for 5 min and the pellet was resuspended in hESC medium (20% knockout serum replacement; Life Technologies), 3% ESC-quality fetal bovine serum (Thermo Fisher Scientific, USA), 0.7% 2-mercapto-ethanol, 1% minimum essential medium non-essential amino acids (MEM-NEAAs; Life Technologies), 1% GlutaMAX (Life Technologies, Canada), and 1% penicillin-streptomycin (P/S; Life Technologies).

For embryoid body (EB) formation, cells were plated at a density of 9,000 cells per well in ultra-low attachment 96-well plate in hESC medium supplemented with 4 ng/ml b-FGF and 50 μM ROCKi. Plate was centrifuged at 300 rpm for 1 min and the medium changed at day-3 and day-5. At day-7, EBs were transferred to ultra-low attachment 24 well plates and neuroinduction medium was added [1% N2 supplement (Gibco), 1% GlutaMAX (Life Technologies), 1% MEM-NEAAs, 1% P/S, and 1 μg/ml heparin in DMEM/F12 (Life Technologies)]. After two days, neuroinduction media was changed and at day-9 EBs were embedded in Matrigel for 1 hour. EBs were placed back into ultra-low attachment 24-well plates and media was changed to differentiation medium minus vitamin A (50% neurobasal medium, 0.5% N2, 1% B27 supplement without vitamin A, 1:100 2-mercapto-ethanol, 0.5% MEM-NEAA, 1% GlutaMAX, and 1:100 P/S in DMEM/F12). Media was replaced at day-13 and at day-15 brain organoids were transferred to 6 well plates under agitation at 90 rpm. At this time point, media was changed to differentiation media plus vitamin A (50% neurobasal medium, 0.5% N2, 1% B27 supplement with vitamin A, 1:100 2-mercapto-ethanol, 0.5% MEM-NEAA, 1% GlutaMAX, and 1:100 P/S in DMEM/F12) and replaced every 4 days until day-45.

### Sample preparation and Liquid-chromatography mass spectrometry (LC-MS)

Brain organoids at day-45 were treated with 7.5 μM harmine or vehicle (DMSO) for 24 hours. At least five organoids per condition from three independent batches were collected and stored at −80°C prior to analysis. Organoids were lysed in a buffer containing 7M Urea, 2M thiourea, 1% CHAPS, 70 mM DTT, and Protease Inhibitor Cocktail (Roche). Supernatant protein extracts (50μg) were digested in gel with trypsin (1:100, w/w) overnight. Peptides were injected into a reverse-phase two-dimensional liquid chromatographer [Acquity UPLC M-Class System (Waters Corporation, USA)], coupled to a Synapt G2-Si mass spectrometer (Waters Corporation, USA). Data was acquired with Data-Independent Acquisitions (DIA) and with ion mobility separation (high-definition data-independent mass spectrometry; HDMSE)^72^. Peptides (5μg) were loaded onto an M-Class BEH C18 Column (130 Å, 5μm, 300μm X 50 mm, Waters Corporation, Milford, MA) in first-dimension chromatography. Three discontinuous fractionation steps were performed with ascending acetonitrile concentrations (13%, 18%, and 50%). After each step, peptide loads were directed to second-dimension chromatography on a nanoACQUITY UPLC HSS T3 Column (1.8μm, 75μm X 150 mm; Waters Corporation, USA). Peptides were then eluted using a 7 to 40% (v/v) acetonitrile gradient for 54 min at a flow rate of 0.4μL/min directly into a Synapt G2-Si. MS/MS analyses were performed by nano-electrospray ionization in positive ion mode nanoESI (+) and a NanoLock Spray (Waters Corporation, Manchester, UK) ionization source. The lock mass channel was sampled every 30 seconds. Mass spectrometer was calibrated with a spectrum reference with [Glu1]-Fibrinopeptide B human (Glu-Fib) solution, using the NanoLock Spray source. All samples were run in technical duplicates of three biological replicates.

### Database search and quantification

Raw files were processed for identification and quantification using *Progenesis^®^ QI for proteomics version 4.0* including software package with peptide 3D, Apex3D, and ion accounting informatics (Waters). Ion accounting and quantitation were performed by searching algorithms and cross-matching with the Uniprot human proteome database, version 2018/09, with the default parameters. Reversed database queries were appended to the original database in order to assess the false-positive identification rate. Protein and peptide level false discovery rate (FDR) was set at 1%. A limit of one missed cleavage was allowed due to digestion by trypsin, and carbamidomethylation was considered a fixed modification while methionine oxidation was considered a variable modification. Identifications that did not satisfy these criteria were rejected. Once LC-MS data is loaded to the software, it performs alignment and peak detection, creating a list of interesting peptide ions (peptides). This list is explored within Peptide Ion Stats by multivariate statistical methods and the final step is protein identification. Relative quantitation of proteins used the Hi-N (3) method of comparison of peptides.

### *In silico* analysis

The normalized intensity data with the identified proteins list was performed using the Perseus software. Fold change of the intensities assigned to each protein was calculated within each experimental batch. Statistical analyses were performed with one sample t-test (p<0.05) against 0 value (no change). The final dataset of differentially expressed proteins was uploaded to functional enrichment analysis at different databases. Gene ontology was analyzed using Panther database^73^. The significantly enriched processes are based on Fisher’s exact test. To identify overrepresented pathways, we used Reactome Pathways Knowledgebase applying a p-value cutoff ≤ 0.05^74^ Protein interaction networks were identified using the STRING database considering medium (0.4) and high (0.7) confidence interactions^75^.

### Immunocytochemistry

Brain organoids were treated with 7.5 μM harmine for 24 hours and fixed overnight in 4% paraformaldehyde (Sigma-Aldrich, USA). Organoids were incubated in 30% sucrose for dehydration overnight at 4°C. Organoids were then embedded in optimal cutting temperature compound (OCT) and stored at −80°C. The organoids were sectioned with a cryostat (Leica Biosystems, Germany) into 20 μm thickness. For immunofluorescence, slides were incubated at 37°C for 30 min and washed in 1X PBS and permeabilized in 0.3% Triton X-100 in PBS for 15 min. For cleaved caspase-3 staining, antigen retrieval procedure was performed before the permeabilization step. Sections were incubated in 10mM citrate buffer, 0.05% Tween 20, pH=6 for 10 min at 98°C. Following this step, slides were incubated in blocking solution containing 1% BSA + 10% normal goat serum (NGS) in PSB for 2 hours. Primary antibodies (see **Supplementary Table 1**) were diluted in blocking solution and incubated at 4°C overnight. Slides were washed in PBS 3 times for 5 min and incubated with re-blocking solution containing 1% BSA + 10% normal goat serum (NGS) in PSB for 20 minutes. After the re-blocking step, slides were incubated with secondary antibodies (1:400 dilution in 1.5% BSA in PBS) for 2 hours at room temperature and then washed 3 times for 5 min in PBS. For nuclear staining, sections were incubated with 300nM DAPI for 10 min. Slides were washed again 3 times for 5 min in PBS and then cover-slipped with Aqua-Poly/Mount (Polysciences, Inc). Images were acquired using a Leica TCS SP8 confocal microscope.

For quantification, tilescan images from 4-5 organoids per condition (vehicle and harmine) were analyzed using Image J software. For each slice, threshold for DAPI and cleaved-caspase 3 were set and the percentage of stained area was used as a parameter to estimate cleaved-caspase 3 expression.

### Western Blot

Organoids were incubated with sample buffer without bromophenol blue (62.5 mM Tris-HCl, pH 6.8, containing 10% glycerol, 2% SDS and 5% 2-mercaptoethanol), vortex thoroughly and homogenized with a polypropylene pestle. Tissue extracts were boiled at 95°C for 5 min, centrifuged at 4°C 15000 g for 15 min and supernatant was collected. Total protein concentration was estimated using the Bio-Rad Protein Assay (#5000006, Biorad). After addition of bromophenol blue (0.02%), extract samples were separated by electrophoresis on a 10% SDS polyacrylamide gel and transferred to polyvinylidene difluoride (PVDF) membranes.

Membranes were blocked in 5% non-fat milk diluted in Tris-Buffered Saline with 0.1% Tween-20 (TBS-T) for 1 hour at room temperature and incubated overnight at 4°C with primary antibodies diluted in TBS-T with 5% non-fat milk. Membranes were then washed with TBS-T and incubated with peroxidase-conjugated antibodies (see **Supplementary Table 1**). Protein bands were visualized using ECL Prime Western Blotting System (#GERPN2232, Amersham) for five minutes and chemiluminescence was detected with an Odyssey-FC System^®^ (Imaging System - LI-COR Biosciences).

To re-use membranes, stripping protocol was performed by incubation with stripping buffer (pH 2.2, 200 mM glycine, SDS 0,1% and 1% Tween-20) for three cycles of 10 minutes. Membranes were then washed with TBS 1X and TBS-T for two cycles of 10 minutes in each solution. Next, membranes were blocked and incubated again with primary antibodies. Densitometry analysis was performed using ImageJ software and the values obtained represent the fold change of the ratio between the immunodetected protein and the loading control (actin).

### Total RNA isolation and PCR

Total RNA was isolated from at least five organoids per batch using PureLink™ RNA Mini Kit (Invitrogen, 12183018A) following the supplier’s instructions. The amount of RNA was evaluated using NanoDrop 2000 (Thermo Fisher Scientific™, ND2000USCAN). Each sample was digested with 4 U of DNAse I Amplification Grade (Invitrogen, 18068-015) and the reverse transcription for cDNA synthesis was performed using M-MLV (Invitrogen, 28025-013). The amplification reactions for qualitative PCR were performed using Recombinant Taq DNA polymerase (Invitrogen, 11615-038) according to the following parameters: 5 minutes at 95°C (denaturation step); 40 cycles of 10 seconds at 95°C; annealing step for 10 seconds at respective temperature for each amplicon (**Supplementary Table 2**) and elongation step for 13 seconds at 72°C. Primers sequences are also listed in **Supplementary Table 2**. PCR products were analyzed by gel electrophoresis (1% agarose and 120V for 40 minutes).

RT-qPCR reactions were performed in three replicates with 10 ng of cDNA per reaction at a final reaction volume of 10 μL in MicroAmp™ Optical 96-Well Reaction Plate (Applied Biosystems™, N8010560) containing 1X GoTaq qPCR Master Mix (Promega Corporation), 300 nM CXR Reference Dye and final concentration 200nM of each SYBR green designed primers described in **Supplementary Table 2**. The reactions and analyses were performed in StepOnePlus Real-Time PCR System (Applied Biosystems™, 4376600). The data were analyzed as fold change (2-ΔΔCt) with gene expression normalized by housekeeping genes *(GAPDH* and *HPRT1*).

### Statistical Analysis

Data are expressed as mean ± SEM. Data comprehending three independent experiments with two conditions for comparison were analyzed by paired two-tailed Student’s t-test.

## Acknowledgments

This work is part of the Master thesis of K.K. We thank Ismael Gomes, Gabriela Vitória, and Beatriz Mello for the technical support on brain organoid culture and slides preparation for immunocytochemistry. We also thank Iván Fernandez and Tomás Falzone for scientific discussions and input.

## Author Contributions

K.K. and S.R. conceptualized the study. K.K. wrote the first draft of the manuscript. M.N.C. performed brain organoid culture. M.N.C. and K.K. performed the harmine treatment and sample collection to proteomics. J.M.N. and D.M.S. performed LC-MS and database search and quantification. L.G.S. performed data normalization and statistical analysis of differentially expressed proteins. K.K. performed *in silico* analysis and prepared the figures. K.K. and J.A.S. performed the RT-qPCR analysis. K.K. and I.M.O. performed western blot analysis. All authors reviewed and approved the final version of the manuscript. D.M.S. and S.R. coordinated the study.

