## Supplementary Material for "Proteomic changes induced by harmine in human brain organoids reveal signaling pathways related to neuroprotection"

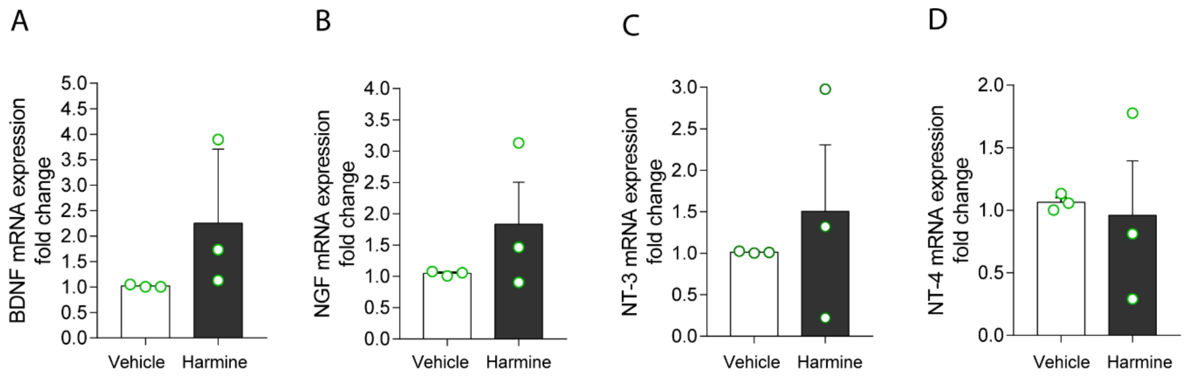


**Supplementary Figure 1:** RT-qPCR for (A) BDNF, (B) NGF, (C) NT-3, and (D) NT-4 in brain organoids treated with harmine for 24 hours. Each circle represents independent experiments. mRNA extraction was performed with at least five organoids per batch.

**Supplementary Table 1: Primary and secondary antibodies used for ICC and WB**

| Antibody | Catalog/Supplier | Dilution |
|---|---|---|
| Rabbit anti-Nestin | RA22125, Neuromics | 1:750 |
| Mouse anti-GFAP | MO15052, Neuromics | 1:100 |
| Mouse anti-MAP2 | M1406, Sigma-Aldrich | 1:300 |
| Rabbit anti-cleaved caspase 3 | AB3623, Merck Millipore | 1:50 |
| Rabbit anti-DYRK1A | ab65220, Abcam | 1:750 |
| Rabbit anti-5HT2A | PA5-103377, Invitrogen | 1:100 |
| Mouse anti-Actin | MAB1501, Merck Millipore | 1:2000 |
| Rabbit anti-CREB | 9197S, Cell Signaling | 1:2000 |
| Rabbit anti p-CREB (Ser133) | 9198S, Cell Signaling | 1:1000 |
| Rabbit anti-AKT (pan) | 4691, Cell Signaling | 1:3000 |
| Rabbit anti p-AKT (Ser473) | 4060, Cell Signaling | 1:1000 |
| Goat anti-Rabbit Alexa Fluor® 488 | A-11008, Invitrogen | 1:400 |
| Goat anti-Mouse Alexa Fluor® 546 | A-11003, Invitrogen | 1:400 |
| Goat anti-Mouse Alexa Fluor® 488 | A-11001, Invitrogen | 1:400 |

| | | |
|---|---|---|
| Goat anti-Rabbit Alexa Fluor® 594 | A-11037, Invitrogen | 1:400 |
| Goat anti-Rabbit HRP conjugate | G21234, Mol. Probes | 1:10000 |
| Goat anti-Mouse HRP conjugate | G21040, Mol. Probes | 1:10000 |

**Supplementary Table 2: Primers sequences and PCR conditions**

| Target gene | Primer sequence | Product (bp) | Annealing point (°C) |
|---|---|---|---|
| 5-HT2a | Forward: ACTCGCCGATGATAACTTTGTCCT<br>Reverse: TGACGGCCATGATGTTTGTGAT | 359 | 58 |
| GAPDH | Forward: TTCGACAGTCAGCCGCATC<br>Reverse: GACTCCACGACGTACTCAGC | 352 | 58 |
| DYRK1A | Forward: GGCTTGCACCGTCATTTTCAT<br>Reverse: GTCACTGTACTGATGTGAATGGG | 80 | 59 |
| BDNF | Forward: AGAGGCTTGACATCATTGGCTG<br>Reverse: CAAAGGCACTTGACTACTGAGCATC | 147 | 57 |
| NGF | Forward: GACTCCGTTCACCCCGTGTGC<br>Reverse: CACACCGAGAATTCGCCCCTG | 166 | 60 |
| NT-3 | Forward: TGGGGGAGACTTTGAATGAC<br>Reverse: CTGGCAAACTCCTTTGATCC | 201 | 55 |
| NT-4 | Forward: AGGAGGCACTGGGTATCTGA<br>Reverse: ATCCCTGAGGTCTCTCAGCA | 198 | 59 |
| HPRT1 | Forward: CGTCGTGATTAGTGATGATGAACC<br>Reverse: AGAGGGCTACAATGTGATGGC | 176 | 60 |
| GAPDH (RT-qPCR) | Forward: CCATCTTCCAGGAGCGAGATC<br>Reverse: TGAAGACGCCAGTGGACTC | 84 | 60 |
